# ACBD3 is an essential pan-enterovirus host factor that mediates the interaction between viral 3A protein and cellular protein PI4KB

**DOI:** 10.1101/375550

**Authors:** Heyrhyoung Lyoo, Hilde M. van der Schaar, Cristina M. Dorobantu, Jeroen R.P.M. Strating, Frank J.M. van Kuppeveld

## Abstract

The enterovirus genus of the picornavirus family includes a large number of important human pathogens such as poliovirus, coxsackievirus, enterovirus-A71, and rhinoviruses. Like all other positive-strand RNA viruses, genome replication of enteroviruses occurs on rearranged membranous structures called replication organelles (ROs). Phosphatidylinositol 4-kinase IIIβ (PI4KB) is required by all enteroviruses for RO formation. The enteroviral 3A protein recruits PI4KB to ROs, but the exact mechanism remains elusive. Here, we investigated the role of Acyl-coenzyme A binding domain containing 3 (ACBD3) in PI4KB recruitment upon enterovirus replication using ACBD3-knockout (ACBD3^KO^) cells. ACBD3 knockout impaired replication of representative viruses from four enterovirus and two rhinovirus species. PI4KB recruitment was not observed in the absence of ACBD3. The lack of ACBD3 also affected the localization of individually expressed 3A, causing 3A to localize to the endoplasmic reticulum instead of the Golgi. Reconstitution of wt ACBD3 restored PI4KB recruitment and 3A localization, while an ACBD3 mutant that cannot bind to PI4KB restored 3A localization, but not virus replication. Consistently, reconstitution of a PI4KB mutant that cannot bind ACBD3 failed to restore virus replication in PI4KB^KO^ cells. Finally, by reconstituting ACBD3 mutants lacking specific domains in ACBD3^KO^ cells, we show that Acyl-coenzyme A binding (ACB) and charged amino acids region (CAR) domains are dispensable for 3A-mediated PI4KB recruitment and efficient enterovirus replication. Altogether, our data provide new insight into the central role of ACBD3 in recruiting PI4KB by enterovirus 3A and reveal the minimal domains of ACBD3 involved in recruiting PI4KB and supporting enterovirus replication.

**Importance:** As all other RNA viruses, enteroviruses reorganize host cellular membranes for efficient genome replication. A host lipid kinase, PI4KB, plays an important role on this membrane rearrangement. The exact mechanism of how enteroviruses recruit PI4KB was unclear. Here, we revealed a role of a Golgi-residing protein, ACBD3, as a mediator of PI4KB recruitment upon enterovirus replication. ACBD3 is responsible for proper localization of enteroviral 3A proteins in host cells which is important for 3A to recruit PI4KB. By testing ACBD3 and PI4KB mutants that abrogate the ACBD3-PI4KB interaction, we showed that this interaction is crucial for enterovirus replication. The importance of specific domains of ACBD3 was evaluated for the first time, and the essential domains for enterovirus replication were identified. Our findings open up a possibility for targeting ACBD3 or its interaction with virus as a novel strategy for a broad-spectrum antiviral drug.

## Introduction

The *Picornaviridae* family is a large group of viruses with a single-stranded, positive-sense RNA genome. Members of the *Enterovirus* genus, which includes poliovirus [PV], coxsackievirus [CV], enterovirus A71 [EV-A71], EV-D68, and rhinovirus [RV], can cause diverse human diseases such as poliomyelitis, meningitis, hand-foot-and-mouth disease, respiratory illness (1). Even though enteroviruses are associated with a variety of clinical manifestations, there are currently no approved vaccines against most enteroviruses except for PV and EV-A71, and antiviral drugs are not available.

All positive-sense RNA viruses, including picornaviruses, induce reorganization of host cellular membranes (2–4) into so called replication organelles (ROs). ROs are enriched with viral replication factors and co-opted host factors, and serve several important purposes in virus replication (5) including facilitating genome replication. Among picornaviruses, enteroviruses and kobuviruses exploit a similar mechanism for RO formation. A host factor, phosphatidylinositol 4-kinase type III beta (PI4KB) is recruited to the replication sites by viral 3A protein (6–8). PI4KB is a cytosolic lipid kinase that must be recruited to membranes to exert its function and there it generates phosphatidylinositol 4-phosphate (PI4P)-enriched environment (7, 9). PI4P recruits and concentrates cellular proteins, and possibly also viral proteins, to facilitate viral genome replication (10, 11). Among the cellular proteins that interact with PI4P are lipid-transfer proteins, such as oxysterol binding protein (OSBP) (12). In normal condition, OSBP creates membrane contact sites between endoplasmic reticulum (ER) and PI4P-enriched trans-Golgi membranes and shuttles cholesterol in exchange for PI4P (13). In a similar manner, OSBP is recruited to RO membranes and mediates a PI4P-dependent flux of cholesterol from ER to ROs (14).

In uninfected cells, PI4KB is recruited to Golgi membranes among others by the small GTPase ADP-ribosylation factor 1 (Arf1) (15) or by acyl-CoA binding domain containing 3 (ACBD3) (7, 8, 16). Kobuviruses recruit PI4KB through ACBD3, which directly interacts with 3A (7, 11). Recently, the crystal structure of the kobuvirus 3A-ACBD3 complex became available, which revealed the binding sites that are important for the 3A-ACBD3 interaction (6). Point mutations in 3A and ACBD3 at the binding interface inhibited the activation of PI4KB (17), suggesting that PI4KB recruitment to membranes via 3A-ACBD3-PI4KB interaction is necessary for kobuviruses to exploit PI4KB activity.

The 3A protein of several enteroviruses (*e.g*., PV and coxsackievirus B3 [CVB3]) binds to brefeldin A resistance guanine nucleotide exchange factor 1 (GBF1), a guanine exchange factor that activates the small GTPase Arf1. Arf1 interacts with PI4KB in non-infected cells. However, evidence has been presented that PI4KB recruitment by CVB3 and RV 3A likely occurs independently of GBF1 and Arf1 (18, 19). A number of enterovirus 3A proteins have been shown to bind to ACBD3 (8, 18). Therefore, several studies have investigated whether enteroviruses depend on ACBD3 to recruit PI4KB. While in one study knockdown of ACBD3 in HeLa cells inhibited poliovirus (PV) replication (8), another reported no inhibition of PV replication in ACBD3-knockdown HEK-293T, IMR5, and HeLa cells (20). In our previous work, we did not observe inhibition of CVB3 or RV replication and no effects on PI4KB recruitment upon ACBD3 knockdown (18, 19).

Here, we re-evaluated the importance of ACBD3 for enterovirus replication using ACBD3 knockout (ACBD3^KO^) cells. We observed that ACBD3 supports replication of representative viruses of different human enterovirus species (EV-A/B/C/D, RV-A/B) by mediating PI4KB recruitment by 3A. For the first time, we showed that the interaction between ACBD3 and PI4KB is crucial for enterovirus replication. In addition, we dissected the different domains of ACBD3 and uncovered that the glutamine-rich region (Q) and Golgi dynamics domain (GOLD) together suffice to support enterovirus replication. Furthermore, our data suggest that ACBD3 is important for proper 3A localization. Overall, our findings implicate that ACBD3 is not just an intermediate through which 3A recruits PI4KB, but may play a central role in RO formation by scaffolding viral proteins and host proteins.

## Results

### ACBD3 knockout inhibits replication of enterovirus A-D and rhinovirus A-B species

Previously, we observed no effects on CVB3 and RV replication in HeLa cells, in which ACBD3 was knocked down for more than 90% (18, 19). Here, we set out to study enterovirus replication in ACBD3^KO^ HeLa cells. HeLa cells lacking ACBD3 were generated with CRISPR-Cas9 technology, and the knockout was confirmed by Western blot analysis (Fig. S1A). Next, we evaluated enterovirus replication kinetics in ACBD3^KO^ cells using representative viruses of four different human enterovirus species (EV-A71 [EV-A], CVB3 [EV-B], PV-1 [EV-C], EV-D68 [EV-D]) and two rhinovirus species (RV-A2 [EV-A] and RV-B14 [RV-B)]). RV-C was not tested, as HeLa R19 cells are unsusceptible to RV-C due to the lack of its receptor, cadherin-related family member 3 (CDHR-3) (21). All of the viruses clearly showed deficient replication in ACBD3^KO^ HeLa cells (Fig. 1A). Replication of enteroviruses was also impaired in another human cell line, haploid human cell line HAP1, in which ACBD3 was knocked out (Fig. S2).

**Figure 1.**
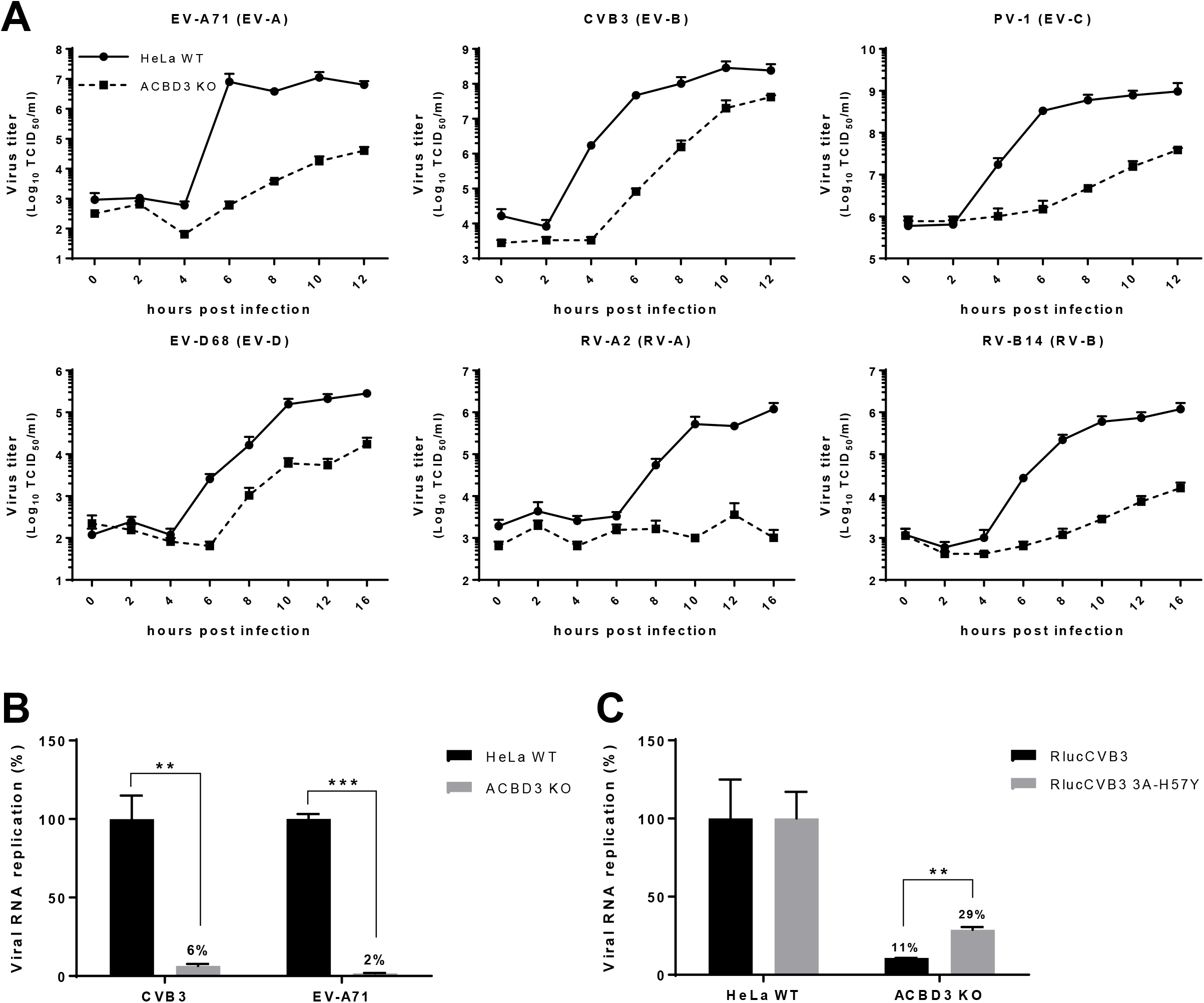
ACBD3 is crucial for enterovirus replication. (A) Growth curves of enteroviruses in HeLa^wt^ and ACBD3^KO^ cells. After infection for 30min at an MOI 5, cells were incubated for the indicated times. Then, cells were freeze-thawed three times to harvest infectious virus particles. Total virus titers were determined by endpoint dilution. (B) RNA replication of CVB3 and EV-A71 virus in HeLa ACBD3^KO^ cells. HeLa^wt^ and ACBD3^KO^ cells were transfected with *in vitro* transcribed RNA of the CVB3 or EV-A71 subgenomic replicons encoding firefly luciferase in place of the capsid region. After 7 h, cells were lysed to determine the intracellular luciferase activity. (C) RNA replication of CVB3 mutant in ACBD3^KO^ cells. HeLa^wt^ and ACBD3^KO^ cells were infected with wt or 3A-H57Y mutant CVB3 reporter viruses carrying a *Renilla* luciferase (RlucCVB3) at an MOI 0.1. After 8 h, cells were lysed to determine luciferase activity. Bars represent the mean of triplicate values ± SEM. Values were statistically evaluated using a two-tailed paired t-test. **, = *P* < 0.01; ***, = *P* < 0.001.

To exclude the role of ACBD3 in virus entry, we assessed viral RNA replication of subgenomic replicons of CVB3 and EV-A71 transfected in HeLa ACBD3^KO^ cells. Replication of both replicons was reduced, which suggests a role for ACBD3 in the genome replication step (Fig. 1B). Next, to test whether ACBD3 functions in the same pathway as PI4KB in enterovirus replication, we employed a mutant virus that is less sensitive to PI4KB inhibition, CVB3 3A-H57Y (22). While the replication of wt CVB3 (RlucCVB3) was impaired in ACBD3^KO^ cells, the replication of RlucCVB3 3A-H57Y was significantly increased (Fig. 1C). Encephalomyelitis virus (EMCV), which belongs to the genus of *Cardiovirus* within *Picornaviridae* family, depends on PI4KA but not on PI4KB for generating ROs (23). The replication of EMCV was not affected in ACBD3^KO^ cells (Fig. S3) suggesting that the inhibition of CVB3 and EV-A71 replication in ACBD3^KO^ cells is connected to the PI4KB pathway. Overall, these results indicate that ACBD3 is an important host factor for enterovirus replication.

### ACBD3 is indispensable for PI4KB recruitment

To determine the importance of ACBD3 for PI4KB recruitment during enterovirus replication, we investigated PI4KB localization in CVB3-infected ACBD3^KO^ cells. Since we observed delayed virus replication in ACBD3^KO^ cells (Fig. 1B), different time points were chosen for HeLa^wt^ cells and ACBD3^KO^ cells to mitigate possible effects of different replication levels on PI4KB recruitment. As previously shown, the CVB3 3A protein colocalized with ACBD3 (Fig. 2A) and PI4KB (Fig. 2B) throughout infection in infected HeLa^wt^ cells, which implies that both ACBD3 and PI4KB localize to CVB3 ROs. PI4KB was more concentrated in 3A-positive cells compared to 3A-negative cells, which suggests that it is actively recruited to virus replication sites. Notwithstanding the similar level of 3A expression compared to HeLa^wt^ cells (Fig. 2A-B), no recruitment of PI4KB was observed in infected ACBD3^KO^ cells at any time point (Fig. 2D). These results indicate that ACBD3 mediates recruitment of PI4KB during enterovirus replication.

**Figure 2.**
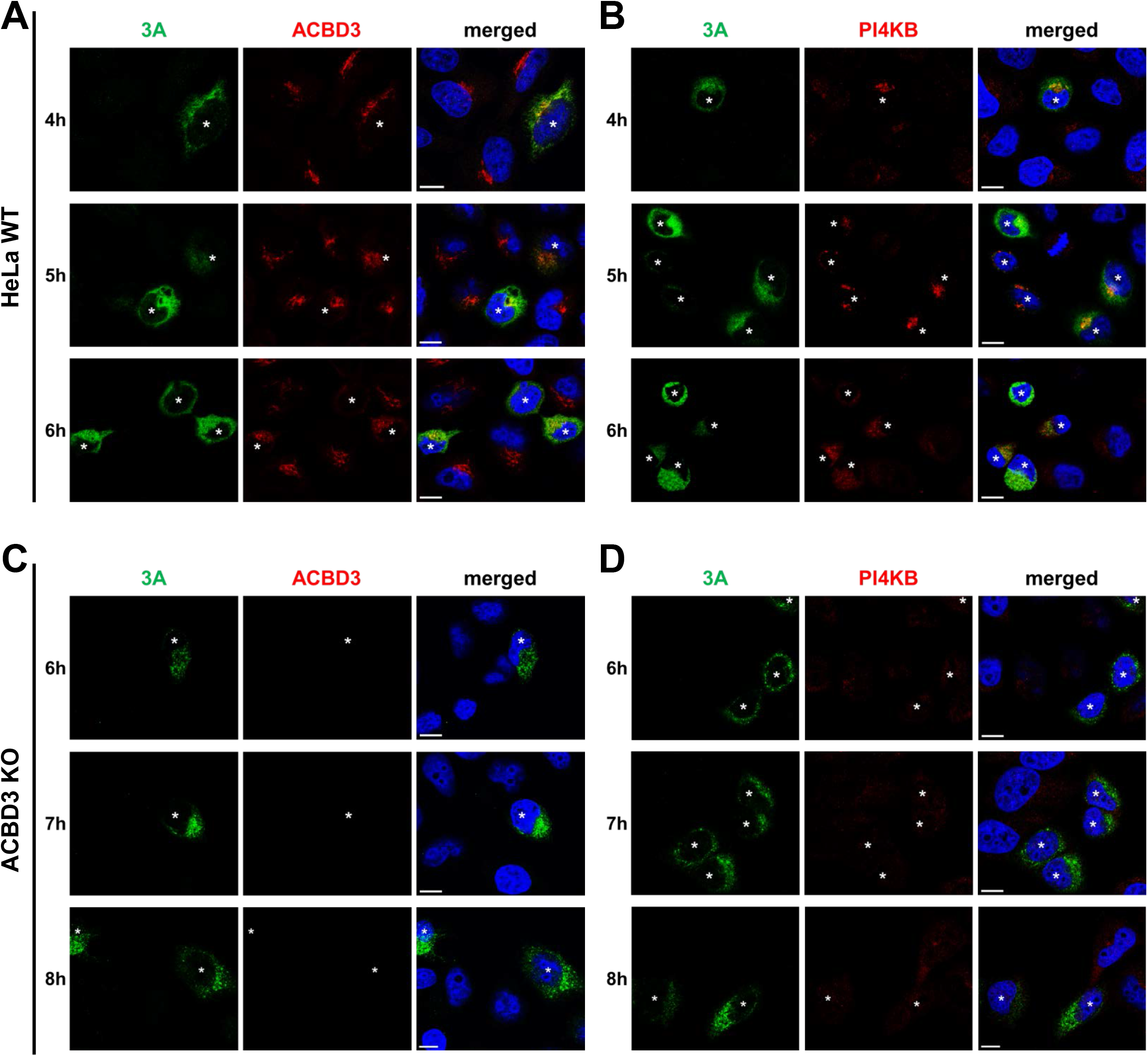
PI4KB recruitment to virus replication sites depends on ACBD3. (A and B) HeLa^wt^ and (C and D) ACBD3^KO^ cells were infected with CVB3 wt at an MOI of 5. At indicated time points, cells were fixed and stained with antibodies against CVB3 3A and ACBD3 (A and C) or CVB3 3A and PI4KB (B and D). Nuclei were stained with DAPI (blue). Asterisks indicate infected cells. Scale bars represent 10 μm.

Enterovirus 3A expression alone is sufficient to recruit PI4KB to membranes (9, 18, 19). To further investigate whether PI4KB recruitment by 3A is mediated by ACBD3, we transiently expressed the 3A proteins from representative human enteroviruses from seven different species (EV-A/B/C/D, RV-A/B/C) with either a C-terminal myc tag or an N-terminal GFP tag and examined the localization of ACBD3 and PI4KB (Fig. 3 and S3). In HeLa^wt^ cells, all 3A proteins colocalized with ACBD3 (Fig. 3A). PI4KB was more concentrated in cells expressing 3A compared to cells that did not express 3A (Fig. 3B), which indicates that PI4KB is actively recruited by enterovirus 3A proteins. In contrast, no PI4KB recruitment was observed in ACBD3^KO^ cells expressing any of the enterovirus 3A proteins (Fig. 3C). These results imply that all enteroviruses utilize a shared mechanism to recruit PI4KB to replication sites, which is via a 3A-ACBD3-PI4KB interaction. Interestingly, we noticed that the localization of 3A differs from HeLa^wt^ cells to ACBD3^KO^ cells (Fig. 3 and S5). Unlike the punctate localization in HeLa^wt^ cells (Fig. 3A), 3A proteins were dispersed throughout the cytoplasm in ACBD3^KO^ cells into a more reticular pattern (Fig. 3B and S5), suggesting that ACBD3 is important for proper localization of 3A.

**Figure 3.**
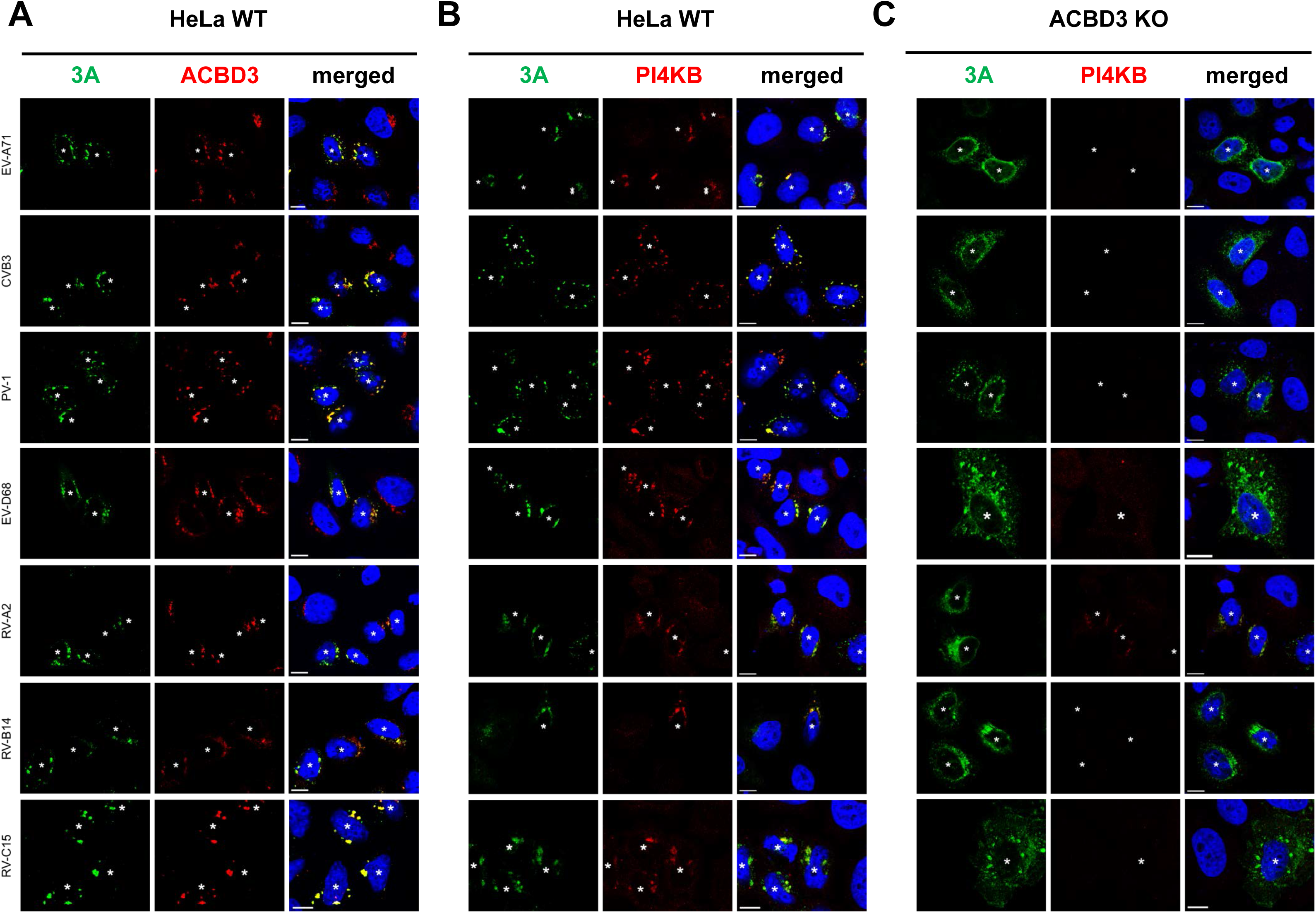
Effects of ACBD3 knockout on the localization of enterovirus 3A proteins and the recruitment of PI4KB. (A) HeLa^wt^ and (B) ACBD3^KO^ cells were transfected with myc-tagged EV-A71 3A, CVB3 3A, PV-1 3A, or EGFP-tagged EV-D68 3A, RV-2 3A, RV-14 3A. The next day, cells were fixed and stained with antibodies against the myc tag to detect 3A, ACBD3, or PI4KB. Asterisks indicate 3A expressing cells. Nuclei were stained with DAPI (blue). Scale bars represent 10 μm.

### ACBD3 is crucial for 3A localization to the Golgi

Because the typical punctate localization of 3A on Golgi-derived membranes is lost in ACBD3^KO^ cells, we assessed the overall structure of the Golgi and ER and the colocalization between 3A and markers for the Golgi (GM130 and TGN46) and ER (Calreticulin). In mock-transfected cells, no gross differences in localization of any of the above markers were observed between HeLa^wt^ cells and ACBD3^KO^ cells (Fig. 4), which suggests that the lack of ACBD3 does not have a major impact on morphology of the Golgi or the ER.

**Figure 4.**
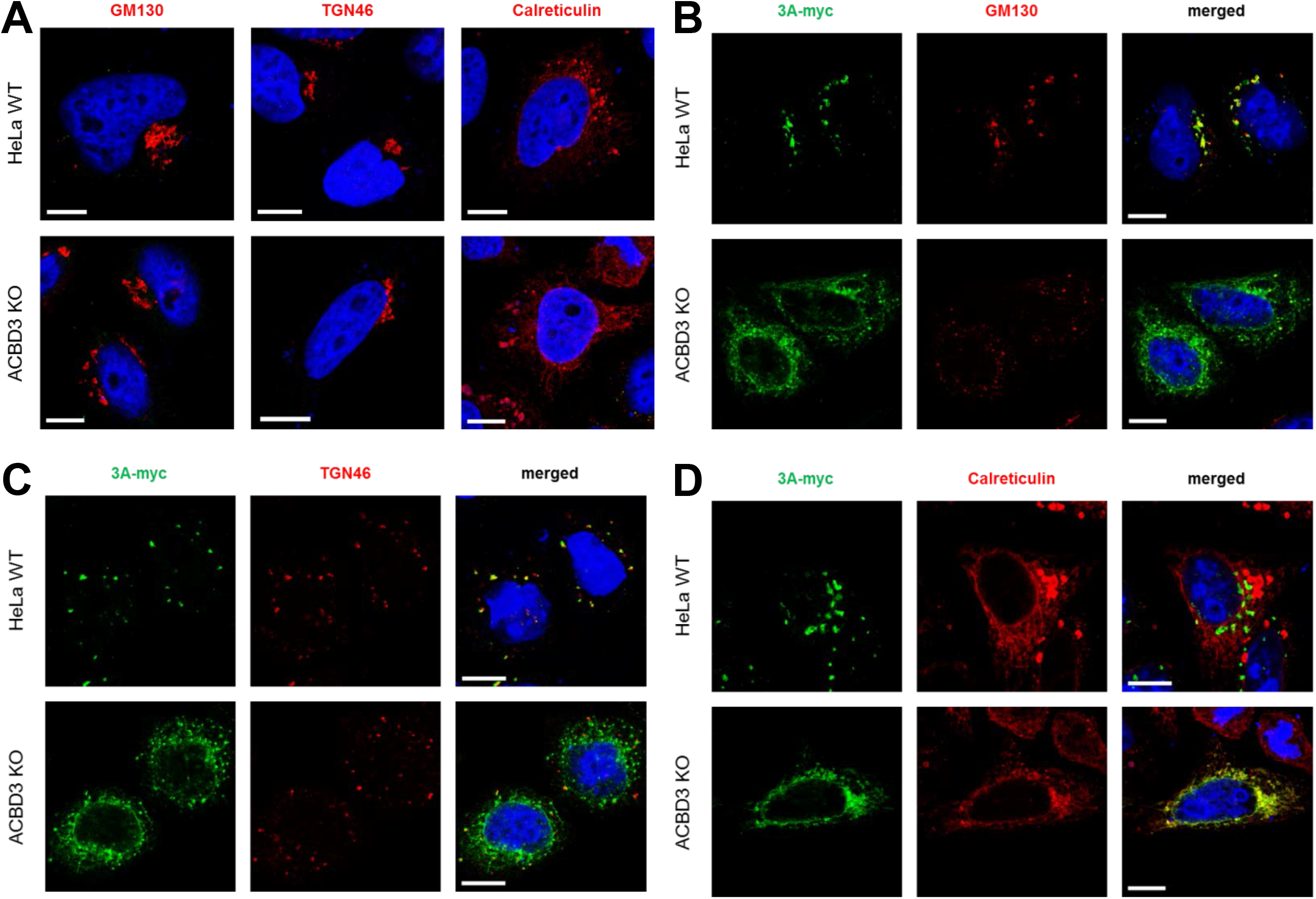
The localization of 3A differs between HeLa^WT^ cells and ACBD3^KO^ cells. (A) Golgi and ER integrity in ACBD3^KO^ cells. HeLa^wt^ and ACBD3^KO^ cells were fixed and stained with antibodies against the Golgi markers GM130 and TGN46 or the ER marker calreticulin. (B-D) HeLa^wt^ and ACBD3^KO^ cells were transfected with myc-tagged CVB3 3A. The next day, cells were fixed and stained with an antibody against the myc tag to detect 3A and with antibodies against GM130 (B), TGN46 (C), or calreticulin (D). Nuclei were stained with DAPI (blue). Scale bars represent 10 μm.

As previously described, the disintegration of the Golgi is likely to be the consequence of the blockage of ER-to-Golgi transport that depends on the interaction between 3A and GBF1/Arf1 (24, 25). In agreement with this, overexpression of 3A caused disassembly of the Golgi apparatus in both wt and ACBD3^KO^ HeLa cells (Fig. 4B-C) pointing out that the disruption of the Golgi by 3A occurs independently of ACBD3. 3A partially colocalized with the Golgi markers but not the ER marker in HeLa^wt^ cells, whereas 3A was localized to the ER, as labeled by calreticulin, in ACBD3^KO^ cells (Fig. 4B-D). This suggests that without ACBD3, 3A cannot localize to the Golgi, which may contribute to the lack of PI4KB recruitment. Collectively, our results suggest that ACBD3 is not merely a mediator between 3A and PI4KB, but plays a central role in recruiting 3A and PI4KB to facilitate virus replication.

### Exogenous expression of wt ACBD3 in ACBD3^KO^ cells restores 3A localization, PI4KB recruitment, and enterovirus replication

To confirm that ACBD3 recruits 3A to the Golgi and mediates the interaction between 3A and PI4KB, we tested whether reconstitution of GFP-tagged ACBD3 in ACBD3^KO^ cells can restore 3A localization and PI4KB recruitment. As a negative control, we used a Golgi localized GFP (coupled to the amino acids 1-60 of galactosyltransferase [GalT]), which failed to restore 3A localization (Fig. 5A; top panel) and PI4KB recruitment (Fig. 5B; top panel). When wt ACBD3 was reconstituted, 3A regained its punctate localization (Fig. 5A; middle panel) and PI4KB was recruited to the same sites (Fig. 5B; middle panel). ACBD3 expressed without 3A was found in the Golgi, where it colocalized with giantin, but no concentrated PI4KB was observed (Fig. S6). These results indicate that proper 3A localization and PI4KB recruitment by 3A depend on ACBD3.

**Figure 5.**
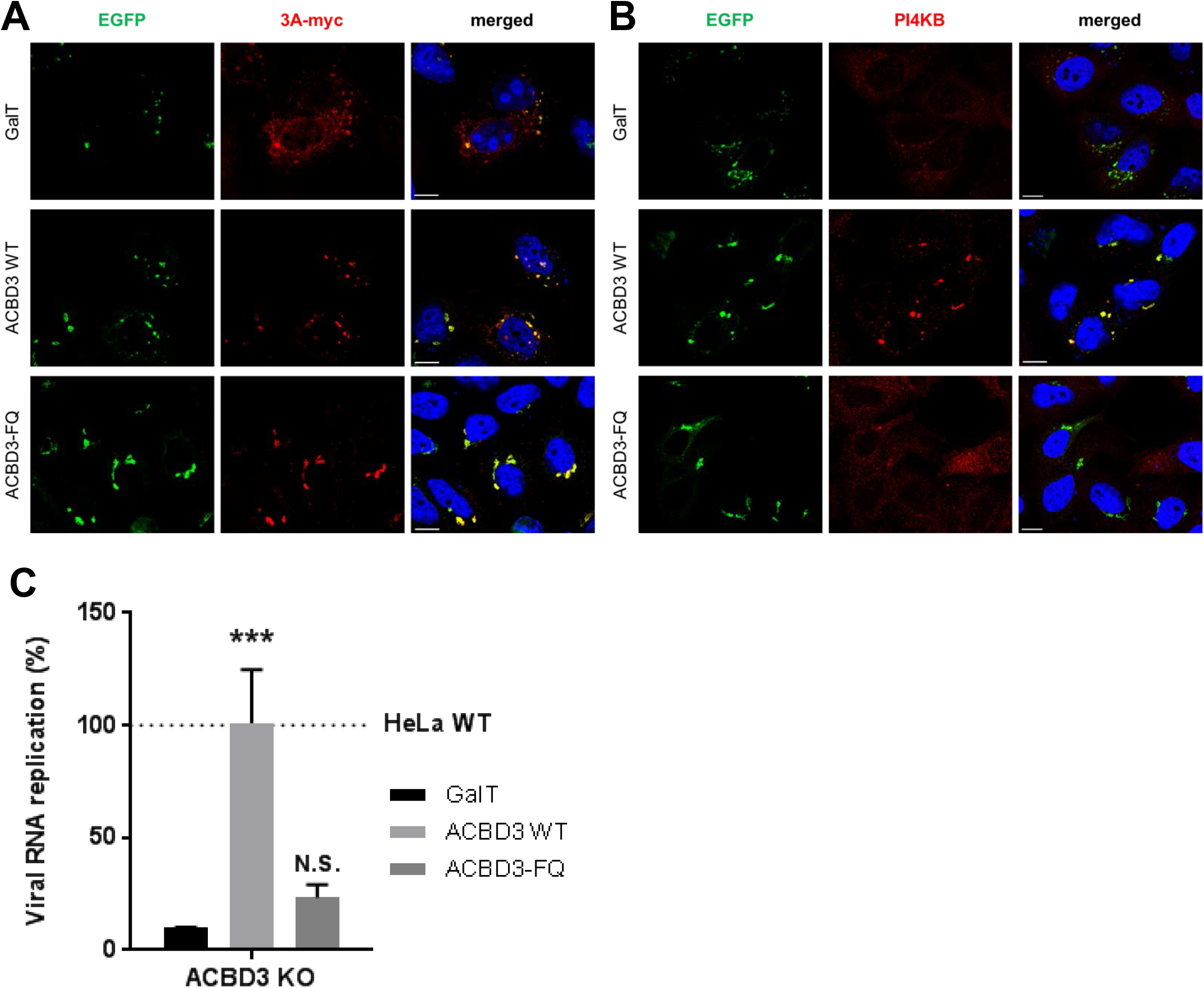
Reconstitution of wt ACBD3 but not ACBD3-FQ mutant rescues PI4KB recruitment and CVB3 replication. (A and B) HeLa ACBD3^KO^ cells were co-transfected with myc-tagged CVB3 3A and EGFP-tagged GalT, ACBD3 wt, or ACBD3-FQ mutant. The next day, cells were fixed and stained with the antibodies against the myc tag to detect 3A (A) or PI4KB (B). Nuclei were stained with DAPI (blue). Scale bars represent 10 μm. (C) HeLa^wt^ and ACBD3^KO^ cells were transfected with EGFP-tagged GalT, ACBD3 wt, or ACBD3-FQ mutant. At 24 h p.t., cells were infected with RlucCVB3 at an MOI of 0.1. After 8 h, cells were lysed to determine luciferase activity. Bars represent the mean of triplicate values ± SEM. Values were statistically evaluated compared to the EGFP control using a one-way ANOVA. ***, = *P* < 0.001; N.S., not significant.

ACBD3 forms a tight complex with PI4KB (16). Recently, it was reported that one or two amino substitution(s) in the Q domain of ACBD3 (F^258^A or F^258^A/Q^259^A) can abrogate binding to PI4KB (16, 17). We employed the F^258^A/Q^259^A mutant (hereafter called “FQ” mutant) to test whether the ACBD3-PI4KB interaction is required for PI4KB recruitment and efficient enterovirus replication. Expression of the ACBD3-FQ mutant restored the punctate localization of 3A (Fig. 5A; bottom panel) but did not support PI4KB recruitment (Fig. 5B; bottom panel). Furthermore, exogenous expression of wt ACBD3 in ACBD3^KO^ cells restored replication of CVB3 to a level comparable to HeLa^wt^ cells, while the negative control (GalT) and ACBD3-FQ mutant could not restore virus replication in ACBD3^KO^ cells (Fig. 5C). Taken together, we showed that 3A localization to the Golgi-derived membranes occurs in an ACBD3-dependent manner and that the interaction between ACBD3-PI4KB is crucial for PI4KB recruitment and efficient virus replication.

### Reconstituted wt PI4KB in PI4KB^KO^ cells can be recruited to the replication sites through the 3A-ACBD3-PI4KB interaction, thereby restoring enterovirus replication

PI4KB is recruited by 3A to ROs during enterovirus replication (8, 9), and depletion of PI4KB by RNAi (9) or pharmacologic inhibition (reviewed in (26)) have been shown to suppress virus replication. Enterovirus mutants resistant to inhibitors of PI4KB contain single amino acid substitutions in the 3A protein (e.g., H57Y for CVB3). CVB3 replication is impaired in PI4KB^KO^ cells that we generated by CRISPR/Cas9 technology (Fig. S1B), while the resistant mutant virus (3A-H57Y) replicated well in PI4KB^KO^ cells (Fig. 6A).

**Figure 6.**
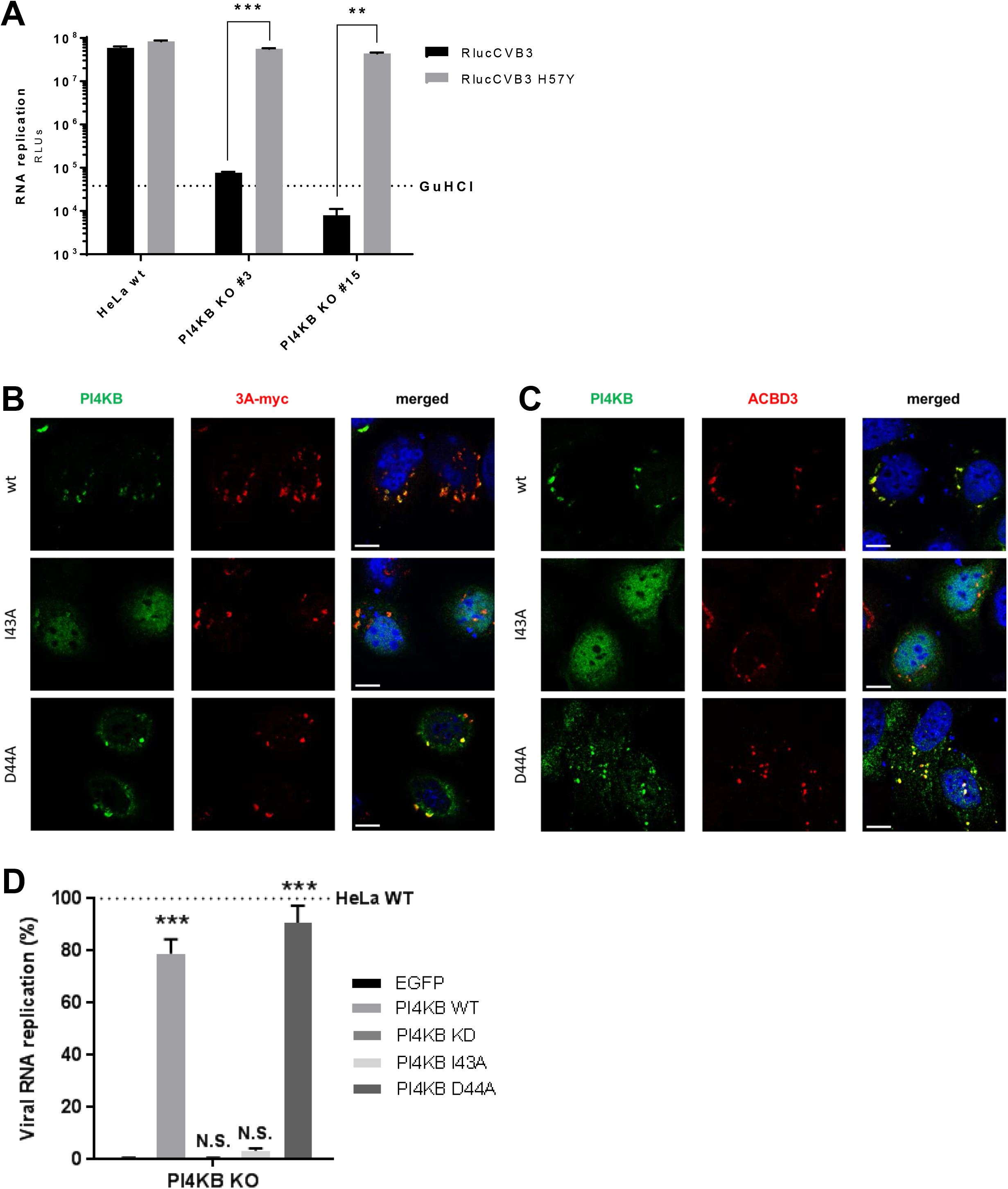
A PI4KB mutant, which does not interact with ACBD3, cannot be recruited by 3A and cannot restore virus replication in PI4KB^KO^ cells. (A) RNA replication of CVB3 mutant in PI4KB^KO^ cells. HeLa^wt^ cells and two PI4KB^KO^ cell clones were infected with wt or 3A-H57Y mutant CVB3 reporter viruses carrying a *Renilla* luciferase (RlucCVB3) at an MOI 0.1. After 8 h, cells were lysed to determine luciferase activity. Bars represent the mean of triplicate values ± SEM. Values were statistically evaluated using a two-tailed paired t-test. **, = *P* < 0.01; ***, = *P* < 0.001. (B, C) HeLa PI4KB^KO^ cells were co-transfected with myc-tagged CVB3 3A and HA-tagged PI4KB wt, PI4KB-I^43^A or D^44^A mutants. The next day, cells were fixed and stained with antibodies against the HA tag to detect PI4KB and against the myc tag to detect 3A (B) or against ACBD3 (C). Nuclei were stained with DAPI (blue). Scale bars represent 10 μm. (D) HeLa^wt^ and PI4KB^KO^ cells were transfected with EGFP, HA-tagged PI4KB wt, PI4KB-I^43^A or D^44^A mutants. At 24 h p.t., cells were infected with RlucCVB3 at an MOI of 0.1. After 8 h, cells were lysed to determine luciferase activity. Bars represent the mean of triplicate values ± SEM. Values were statistically evaluated compared to the EGFP control using a one-way ANOVA. ***, = *P* < 0.001; N.S., not significant.

Two PI4KB mutants (I^43^A and D^44^A) were previously shown to reduce PI4KB binding to ACBD3 *in vitro* (16, 17). While wt PI4KB reconstituted in PI4KB^KO^ cells colocalized with 3A and ACBD3 (Fig. 6B-C; top panel), the PI4KB-I^43^A mutant did not colocalize with 3A and ACBD3, and instead mostly localized to the nucleus (middle panel). Unexpectedly, the D^44^A mutant did colocalize with 3A and ACBD3 (bottom panel). In line with this, wt PI4KB and the D^44^A mutant could restore enterovirus replication in PI4KB^KO^ cells, while the I^43^A mutant and the negative controls, EGFP and a PI4KB kinase-dead mutant that lacks catalytic activity (PI4KB-KD) could not (Fig. 6D). Why the D^44^A mutant behaves differently from the I^43^A mutant is unclear. Possibly, the D^44^A mutant has residual interaction with ACBD3 in cells that could not be detected *in vitro*. Nevertheless, these results imply that the interaction between ACBD3 and PI4KB is important for enterovirus replication.

### Enterovirus replication does not require the ACB and CAR domains of ACBD3

Four domains are recognized in ACBD3; the acyl-CoA binding (ACB) domain, the charged amino acids region (CAR), the glutamine-rich region (Q), and the Golgi dynamics domain (GOLD) (Fig. 7A). The ACB domain, which is relatively conserved among all known ACBD proteins (ACBD1-7), has been suggested to be important for binding to long-chain acyl-CoA (27), and for binding to sterol regulatory element binding protein 1 (SREBP1) causing reduction of *de novo* palmitate synthesis (28). The CAR domain contains a nuclear localization signal (29), yet the function of the CAR domain is unknown. The Q domain interacts with the N-terminal helix of PI4KB (12, 18). The GOLD domain interacts with giantin, and by doing so, tethers ACBD3 to the Golgi membrane (29). Enterovirus and Kobuvirus 3A proteins bind to the GOLD domain, most probably at the same site as giantin (7, 18). To investigate the importance of the ACB and CAR domains for enterovirus replication we tested whether N-terminal deletion mutants of ACBD3 could restore enterovirus replication in ACBD3^KO^ cells. Mut1 and mut2, which contain intact Q and GOLD domains, could restore virus replication to a level comparable to cells reconstituted with wt ACBD3 (Fig. 7B). This is in alignment with our observation that these mutants colocalize with 3A and PI4KB in ACBD3 KO cells (Fig. 7C). In contrast, mut3, which contains only the GOLD domain, could not rescue virus replication (Fig. 7B) or PI4KB recruitment (Fig. 7C), like the negative controls (GalT and the FQ mutant) (Fig. 7B), even though all mutants colocalized with 3A (Fig. 7C). Of note, although mut3 and 3A colocalized in punctate structures, they also partly co-localized to reticular and nuclear envelope-like structures, which may indicate that PI4KB also plays a role in firmly localizing 3A and ACBD3 to Golgi-derived membranes. These results indicate that enterovirus replication requires the Q and GOLD domains of ACBD3 for localization of viral protein 3A to the Golgi and for hijacking PI4KB.

**Figure 7.**
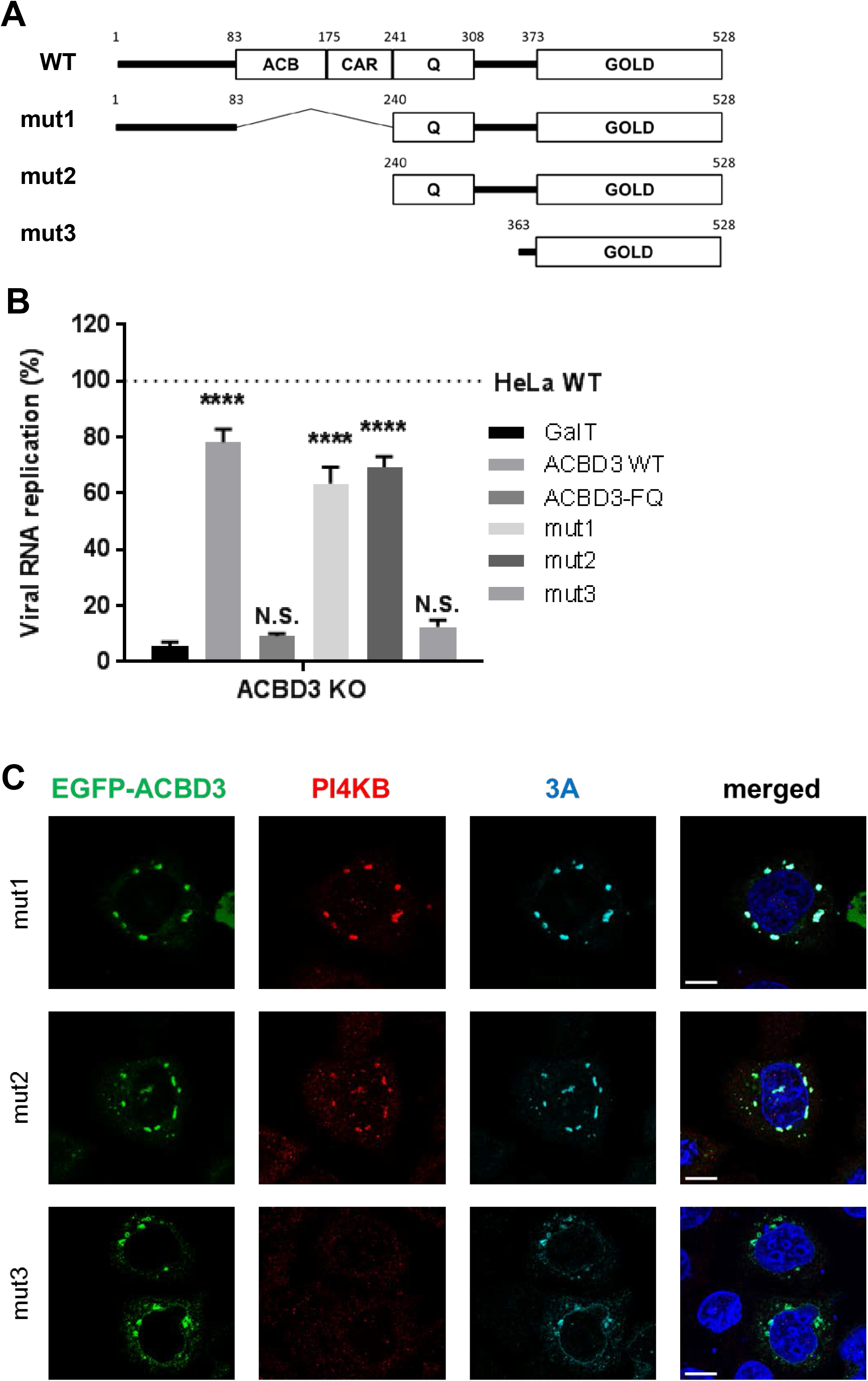
The Q and GOLD domains of ACBD3 are sufficient to support proper 3A localization, PI4KB recruitment, and enterovirus replication. (A) Schematic representation of full-length ACBD3 and its N-terminal deletion mutants (mut1-3). ACBD3 contains the acyl-CoA binding (ACB) domain, the charged amino acids region (CAR), the glutamine rich (Q) domain, and the Golgi dynamics domain (GOLD). Numbers indicate amino-acid positions. (B) HeLa^wt^ and ACBD3^KO^ cells were transfected with EGFP-tagged GalT, ACBD3 wt, ACBD3-FQ mutant, or ACBD3 N-terminal deletion mutants (mut1-3). At 24 h p.t., cells were infected with RlucCVB3 at an MOI of 0.1. After 8 h, cells were lysed to determine luciferase activity. Bars represent the mean of triplicate values ± SEM. Values were statistically evaluated compared to the EGFP control using a one-way ANOVA. ****, = *P* < 0.0001; N.S., not significant. (C) HeLa ACBD3^KO^ cells were co-transfected with myc-tagged CVB3 3A and EGFP-tagged ACBD3 wt, or ACBD3 N-terminal deletion mutants (mut1-3). The next day, cells were fixed and stained with the antibodies against PI4KB (red) and the myc tag to detect 3A (light blue) or. Nuclei were stained with DAPI (blue). Scale bars represent 10 μm.

## Discussion

Both enteroviruses and kobuviruses of the *Picornaviridae* family co-opt PI4KB to build up ROs. Viral protein 3A is responsible for PI4KB recruitment to enterovirus replication sites, yet the underlying mechanism has remained elusive. Despite the direct interaction between ACBD3 and enterovirus 3A proteins (8, 18), there has not yet been a consensus about the importance of ACBD3 for enterovirus replication and PI4KB recruitment. Previously, we observed no inhibition on CVB3 and RV replication and no effects on PI4KB recruitment, even though more than 90% of ACBD3-knockdown was achieved by siRNA (18, 19). In the present study in which we use ACBD3^KO^ cells, we showed that ACBD3 is an important host factor for replication of four different human enterovirus species (EV-A/B/C/D) and two rhinovirus species (RV-A/B). All viruses showed impaired growth in ACBD3^KO^ cells (Fig. 1 and S2). In addition, neither virus infection (Fig. 2) nor the expression of enterovirus 3A proteins alone (Fig. 3) elicited PI4KB recruitment in the absence of ACBD3. In agreement with our data, the inhibition of EV-A71 and CVB3 was recently reported in ACBD3^KO^ cells (30–32). The discrepancy in the role of ACBD3 from KD to KO condition could result from insufficient suppression of ACBD3 function by RNA interference. In fact, this implicates that the small amounts (~10%) of ACBD3 that remained after knockdown is sufficient to support enterovirus replication and PI4KB recruitment. Similar issues on the differences between knockdown and knockout have been raised (33). For instance, the importance of cyclophilin A (CypA) in nidovirus replication was only prominent in CypA knockout cells but not in knockdown condition, even though CypA protein was undetectable after knockdown (34).

We observed that the lack of ACBD3 has a profound effect on enterovirus 3A protein localization. 3A proteins were found almost exclusively at the ER in ACBD3^KO^ cells (Fig. 4D), whereas in HeLa^wt^ cells they showed a punctate localization mostly on Golgi-derived membranes (Fig. 4B-C). Upon reconstitution of wt ACBD3 in ACBD3^KO^ cells, the localization of 3A was restored to a punctate pattern (Fig. 5A). These findings hint at a new role of ACBD3 for enterovirus replication, which is more than merely being a connector between 3A proteins and PI4KB.

Considering that ACBD3 is involved in several different protein complexes, enteroviruses may take advantage of ACBD3 in several ways, more than just for PI4KB recruitment. ACBD3 may be a scaffold responsible for positioning 3A near cellular factors, including other ACBD3-interacting proteins, required for RO formation. For example, ACBD3 and PV1 3A were found in a protein complex together with the putative Rab33 GTPase-activating proteins TBC1D22A/B (35). In addition, several Golgi stacking proteins such as Golgin45 and Golgi reassembly stacking protein 2 (GORASP2) were recently identified as novel interaction partners of ACBD3, and ACBD3 was proposed as a scaffold tethering Golgin45, GRASP55, and TBC1D22 for the formation of a Golgi cisternal adhesion complex at the medial Golgi (36). It is largely unknown which domains of ACBD3 are responsible for the interaction with the above-mentioned interacting partners, and whether these proteins are recruited to enterovirus ROs also remains to be investigated.

The GOLD domain of ACBD3 is responsible for the interaction with enterovirus 3A protein (18, 35), while the Q domain interacts with PI4KB (16, 17). By utilizing mutants of ACBD3 or PI4KB which disturb the interaction with each other (Fig. 5 and 7), we show for the first time that the interaction between ACBD3 and PI4KB is crucial for enterovirus replication. Aside from the Q and GOLD domains, other domains (i.e., ACB and CAR) of ACBD3 seem to be not involved in enterovirus replication (Fig. 7). This indicates that the functions of the ACB and CAR domains, as well as the cellular proteins and/or lipids that interact with these domains, are unlikely required for enterovirus replication. Although we cannot exclude that additional or unidentified proteins that bind to the Q or GOLD domains of ACBD3 might also be important for enterovirus replication, our findings suggest that ACBD3 mainly serves as a mediator of PI4KB recruitment to ROs, involving the Q and GOLD domains.

How exactly enterovirus 3A protein interacts with ACBD3 needs to be further investigated. The GOLD domain of ACBD3 interacts with enterovirus 3A proteins (18). Similarly, kobuvirus 3A interacts with the ACBD3 GOLD domain and the crystal structure of kobuvirus 3A in complex with the ACBD3 GOLD domain was revealed recently (6). According to this structure, kobuvirus 3A wraps ACBD3 and stabilizes ACBD3 on membranes through the membrane-binding features at the myristoylated N-terminal and hydrophobic C-terminal ends of 3A. However, enterovirus 3A proteins can bind to membranes only through the hydrophobic C-terminus, and the 3A proteins of enteroviruses differ greatly in sequence from kobuvirus 3A. Therefore, the way by which enterovirus 3A interacts with ACBD3 could be different from kobuvirus. Thus, structural insight into the enterovirus 3A - ACBD3 GOLD complex is urgently required to understand how enterovirus 3A interacts with ACBD3.

In conclusion, our study reveals that enteroviruses employ a conserved mechanism to recruit PI4KB, which depends on the Golgi-residing protein ACBD3. Furthermore, we suggest that ACBD3 tethers viral and host proteins to form ROs. Considering the pan-enteroviral dependency on ACBD3, targeting ACBD3 or the 3A-ACBD3 interaction presents as a novel strategy for broad-spectrum antiviral drug development.

## Materials and Methods

### Cells and culture conditions

HAP1^wt^ cells and HAP1 ACBD3^KO^ cells were obtained from Horizon Discovery. HeLa R19 cells were obtained from G. Belov (University of Maryland and Virginia-Maryland Regional College of Veterinary Medicine, US). HAP1 cells were cultured in IMDM (ThermoFisher Scientific) supplemented with 10% fetal calf serum (FCS) and penicillin–streptomycin. HeLa cells were cultured in DMEM (Lonza) supplemented with 10% FCS and penicillin–streptomycin. All cells were grown at 37°C in 5% CO_2_.

### Viruses

The following enteroviruses were used: EV-A71 (strain BrCr, obtained from the National Institute for Public Health and Environment; RIVM, The Netherlands), CVB3 (strain Nancy, obtained by transfection of the infectious clone p53CB3/T7 as described previously (37)), RlucCVB3, RlucCVB3 3A-H57Y (obtained by transfection of infectious clones pRLuc-53CB3/T7 as described previously (22)), RlucEMCV (strain Mengovirus, obtained by transfection of the infectious clone pRLuc-QG-M16.1 as described previously (38)), PV1 (strain Sabin, ATCC), EV-D68 (strain Fermon, obtained from RIVM, The Netherlands), RV-2 and RV-14 (obtained from Joachim Seipelt, Medical University of Vienna, Austria). Virus titers were determined by end-point titration analysis and expressed as 50% tissue culture infectious dose (TCID50).

### Virus infection

Virus infections were carried out by incubating subconfluent HAP1 or HeLa cells for 30 min with virus. Following virus removal, fresh medium or medium containing the control inhibitors guanidine hydrochloride (2 mM) or dipyridamole (100 μM) was added to the cells. To determine one-step growth kinetics for each virus, infected cells were frozen from 2 to 16 h post infection (p.i.). Virus titers were determined by end-point titration analysis and expressed as 50% tissue culture infectious dose (TCID_50_). To check for the recruitment of PI4KB upon virus replication, cells were fixed for immunofluorescence staining as described in below section separately. To check genome replication by measuring intracellular *Renilla* luciferase activity, cells were lysed at 8 h p.i. and followed the manufacturer’s protocol (*Renilla* luciferase assay system; Promega).

### RNA transfection

The subgenomic replicons of CVB3 (10) and EV-A71 (39) were described previously. HeLa cells were transfected with RNA transcripts of replicon constructs. After 7 h, cells were lysed to determine intracellular firefly luciferase activity.

### Plasmids

p3A(CVB3)-myc (25), pEGFP-3A(RV-2), and pEGFP-3A(RV-14) were described previously (19). p3A(EV-A71)-myc, p3A(PV1)-myc, pEGFP-3A(EV-D68), and pEGFP-3A(RV-C15) were prepared by cloning cDNA encoding EV-A71 and PV1 3A into p3A(CVB3)-myc vectors from which CVB3 3A was excised using restriction enzyme sites SalI and BamHI, and EV-D68 and RV-C15 3A into pEGFP vectors using restriction enzyme sites BglII and BamHI. pEGFP-GalT was a gift from Jennifer Lippincott-Schwartz (Addgene plasmid #11929). pEGFP-ACBD3 was a gift from Carolyn E. Machamer (Johns Hopkins University, USA). pEGFP-ACBD3-FQ and pEGFP-ACBD3-mut1/mut2/mut3 were generated by using Q5 Site-Directed Mutagenesis kit (New England BioLabs). pCDNA3-PI4KB(wt)-HA was a gift from Tamas Balla (NIH, USA). pCDNA3-PI4KB(D671A) (KD: kinase activity dead mutant), pCDNA3-PI4KB(I43A), and pCDNA3-PI4KB(D44A) were generated by using Q5 Site-Directed Mutagenesis kit (New England BioLabs).

### Replication rescue assay

HeLa cells were transfected with plasmids carrying wt or mutant ACBD3 (FQ, mut1, mut2, mut3), wt or mutant PI4KB (I43A, D44A), Golgi-targeting EGFP (pEGFP-GalT) or kinase-dead PI4KB (PI4KB-KD) as a negative control. At 24 h post transfection, the cells were infected with CVB3-Rluc. At 8 h p.i., the intracellular *Renilla* luciferase activity was determined by using the *Renilla* luciferase assay system (Promega).

### Antibodies

The rabbit antiserum and the mouse monoclonal antibody against CVB3 3A were described previously (18, 24). Mouse monoclonal antibodies included anti-ACBD3 (Sigma), anti-myc (Sigma), anti-GM130 (BD Biosciences), anti-Giantin (a gift from Marjolein Kikkert, Leiden University Medical Center, The Netherlands). Rabbit polyclonal antibodies included anti-PI4KB (Millipore), anti-myc (Thermo Scientific), anti-TGN46 (Novus Biologicals), anti-Calreticulin (Sigma), anti-HA (Santa Cruz). Conjugated goat antirabbit and goat anti-mouse Alexa Fluor 488, 596, or 647 (Molecular Probes) were used as secondary antibodies for immunofluorescence analysis. For Western Blot analysis, IRDye goat anti-mouse or antirabbit (LI-COR) were used.

### Immunofluorescence microscopy

HeLa cells were grown on coverslips in 24-well plates. Subconfluent cells were transfected with 200 ng of plasmids using Lipofectamine 2000 (Thermo) according to the manufacturer’s protocol or infected with CVB3 at an MOI of 5. At 16 h post transfection (p.t.) or 5-9 h p.i., cells were fixed with 4% paraformaldehyde for 15 min at room temperature. After permeabilization with 0.1% Triton X-100 in PBS for 5 min, cells were incubated with primary and secondary antibodies diluted in 2% normal goat serum in PBS. Nuclei were stained with DAPI. Coverslips were mounted with FluorSave (Calbiochem), and confocal imaging was performed with a Leica SpeII confocal microscope.

### Western blot analysis

HAP1 and HeLa cells were harvested and lysed by TEN-lysis buffer (50 mM TrisHCl pH 7.4, 150 mM NaCl, 1 mM EDTA, 1% NP-40, 0.05% SDS). After 30 min incubation on ice, lysates were centrifuged for 20 min at 10,000 xg. Supernatants were boiled in Laemmli sample buffer for 5 min at 95°C. Samples were run on polyacrylamide gels and transferred to a PVDF membrane (Bio-Rad). Membranes were sequentially incubated with primary antibody against ACBD3 or PI4KB at 4°C overnight and secondary antibodies against mouse IgG or rabbit IgG for 1h at room temperature. Images were acquired with an Odyssey imaging system (LI-COR).

## Acknowledgments

This work was supported by research grants from the Netherlands Organisation for Scientific Research (NWO-VENI-863.12.005 to HMvdS, NWO-VENI-722.012.066 to JRPMS, NWO-VICI-91812628, NWO-ECHO-711.017.002 and ERASysApp project ‘SysVirDrug’ ALW project number 832.14.003 to FJMvK) and from the European Union (Horizon 2020 Marie Skłodowska-Curie ETN ‘EUVIRNA’, grant agreement number 264286 and ‘ANTIVIRALS’, grant agreement number 642434 to FJMvK).

